# Constitutive expression of Cas9 and rapamycin-inducible Cre recombinase facilitates conditional genome editing in *Plasmodium berghei*

**DOI:** 10.1101/2024.10.14.618196

**Authors:** Samhita Das, Tanaya Unhale, Carine Marinach, Belsy del Carmen Valeriano Alegria, Camille Roux, Hélène Madry, Badreddine Mohand Oumoussa, Rogerio Amino, Shiroh Iwanaga, Sylvie Briquet, Olivier Silvie

## Abstract

Malaria is caused by protozoan parasites of the genus *Plasmodium* and remains a global health concern. The parasite has a highly adaptable life cycle comprising successive rounds of asexual replication in a vertebrate host and sexual maturation in the mosquito vector *Anopheles*. Genetic manipulation of the parasite has been instrumental for deciphering the function of *Plasmodium* genes. Conventional reverse genetic tools cannot be used to study essential genes of the asexual blood stages, thereby necessitating the development of conditional strategies. Among various such strategies, the rapamycin-inducible dimerisable Cre (DiCre) recombinase system emerged as a powerful approach for conditional editing of essential genes in human-infecting *P. falciparum* and in the rodent malaria model parasite *P. berghei*. We previously generated a DiCre-expressing *P. berghei* line and validated it by conditionally deleting several essential asexual stage genes, revealing their important role also in sporozoites. The advent of CRISPR enabled targeted genome editing with higher accuracy and specificity and greatly advanced genome engineering in *Plasmodium* spp. Here, we developed new *P. berghei* parasite lines by integrating the DiCre cassette and a fluorescent marker in parasites constitutively expressing Cas9. Owing to the dual integration of CRISPR-Cas9 and DiCre, these new lines allow unparalleled levels of gene modification and conditional regulation simultaneously. To illustrate the versatility of this new tool, we conditionally knocked-out the essential gene encoding the claudin-like apicomplexan micronemal protein (CLAMP) in *P. berghei* and confirm the role of CLAMP during invasion of erythrocytes.

## Introduction

Malaria is a life-threatening disease with a majority of the world population at risk [1]. The causative organism, *Plasmodium spp*, spreads the disease through the bite of an infected *Anopheles* mosquito. The bite causes the release of sporozoites into the mammalian skin where the parasite crosses through different cellular layers and travels to the liver. Inside the liver, sporozoites traverse the hepatocytes, before finally invading them by forming a parasitophorous vacuole (PV) within which they further develop into erythrocyte-invasive merozoites. Some merozoites replicate inside erythrocytes establishing the infection, while others develop into gametocytes, which are taken up by the mosquito during a bite, where the parasite develops into sporozoites that invade the salivary glands for transmission to a new mammalian host [2].

The various stages in the life cycle of *Plasmodium spp* are being studied to identify essential cellular processes, molecular targets and vaccine antigens. *P. berghei* is a widely studied rodent malaria model parasite, owing to the physiological similarities with human malaria and ease in laboratory manipulations. Genome editing tools have significantly improved our knowledge on the role of parasite genes, however, functions of many protein-coding genes are yet to be revealed. Transfection of *Plasmodium* parasites is typically performed in asexual blood stages, precluding loss-of-function mutations of genes that are essential in this part of the life cycle. In recent years, gene manipulation strategies have evolved allowing researchers to conditionally control gene expression [3]. The dimerisable Cre (DiCre) recombinase system emerged as a powerful tool for conditional gene deletion, first established in *Toxoplasma gondii* [4] in the Apicomplexan phylum and then adapted in *P. falciparum* [5,6] and *P. berghei* [7]. The DiCre system functions via expression of the Cre recombinase, split in two separate, inactive polypeptides, each fused to a different rapamycin-binding protein. Rapamycin-induced heterodimerization of the two components in the presence of rapamycin restores recombinase activity. The active Cre recognises short 34 bp sequences called *lox* sites, allowing excision of the floxed (flanked by *lox*) DNA sequences with very high efficiency. Along with the efficiency of Cre recombinase, the introduction of *lox* sites is key to efficient genome editing. *Lox* sites can be introduced within the ORF as an artificial intron [5] or within endogenous introns or UTRs. We previously generated a *P. berghei* line expressing the DiCre cassette and illustrated how this tool allows the study of essential genes not only in asexual blood stages, but also in other stages of the parasite life cycle [7]. Conditional gene deletion in PbDiCre parasites revealed the essential role of *ama1*, *ron2*, *ron4* and *clamp* genes in blood stages and sporozoites [8,9]. In these studies, a two-step strategy was employed to insert *lox* sites upstream and downstream of the target gene.

The past decade also observed a novel emergence of versatile clustered regularly interspaced short palindromic repeats (CRISPR)/CRISPR-associated (Cas) nucleases and their variants with increased genome editing efficiency and specificity. The system consists of a Cas9 nuclease which forms a complex with a single-guide RNA (sgRNA) to precisely generate a double strand break (DSB) in the genome. As *Plasmodium* lacks the classical non-homologous end-joining (NHEJ) pathway, DSB can only be repaired by homology-directed recombination (HDR) with a provided donor DNA template [10]. Since its implementation, there has been a lot of technical developments in the use of CRISPR/Cas9 in *Plasmodium spp*. The latest improvements of the CRISPR/Cas9 system in *P. berghei* involves co-transfection of parasites constitutively expressing Cas9 from an integrated cassette with a linearized donor template together with a plasmid encoding one or two sgRNA [11]. This approach was shown to allow genome editing with very high efficiency, from gene knockout to targeted mutagenesis [11].

Here, we aimed to combine both the CRISPR and DiCre strategies and generated *P. berghei* parasite lines harboring Cas9 and DiCre cassettes integrated in their genome together with a fluorescent marker. Our goal was to produce a useful, efficient and versatile tool allowing a wide range of genetic modifications in *P. berghei*, from targeted mutagenesis to conditional genome editing. For this purpose, we used *P. berghei* PbCas parasites containing a Cas9 cassette integrated in the dispensable *p230p* locus cassette under the control of the *HSP70* promoter for constitutive expression [12]. We modified this line using CRISPR to further integrate at the same locus a DiCre cassette and a fluorescent marker (either GFP or mCherry) to facilitate parasite detection. We generated two lines and named them *PbCasDiCre-GFP and PbCasDiCre-mCherry*, containing the GFP or the mCherry fluorescent marker, respectively. To illustrate the versatility and efficiency of this new tool, we used CRISPR to modify the essential gene *clamp* in one single step, and show that rapamycin-induced conditional deletion of *clamp* impairs merozoite invasion.

## Results

### Generation of fluorescent transgenic *P. berghei* lines expressing both Cas9 and dimerisable Cre

Constitutive expression of Cas9 allows genome editing in *P. berghei* with high efficiency following transfection of sgRNA and a linear donor template for DNA repair by homologous recombination [11]. In order to combine the Cas9 and DiCre systems, we used CRISPR to genetically modify PbCas9 parasites that constitutively express Cas9 from a cassette integrated at the dispensable *p230p* gene locus [11,12]. A 20-base sgRNA target was selected in the *p230p* locus, downstream of the integrated Cas9 cassette, and the corresponding sequence was cloned downstream of PbU6 promoter into a sgRNA plasmid containing a pyrimethamine-resistance cassette (**Figure 1A**). For the repair DNA template, we used the same DiCre cassette as previously described [7]. This cassette consists in the N-terminal Cre 59 (residues Thr19-Asn59) and C-terminal Cre 60 (Asn60-Asp343) portions of the Cre fused at their N-terminus to FKBP12 and FRB, respectively. These two components were placed under control of the constitutive bidirectional promoter eEF1alpha, and followed by the 3’ untranslated region (UTR) from PfCAM and PbHSP70, respectively [7]. In this construct, we further incorporated a fluorescent marker (GFP or mCherry, respectively) under the control of the *hsp70* promoter and the 3’ UTR of PbDHFR. On each side of the construct, we inserted 5’ and 3’ homology fragments for integration at the *p230p* locus, downstream of the Cas9 cassette (**Figure 1A**). The donor template DNA was assembled in a plasmid and linearized prior to transfection.

**Figure 1.**
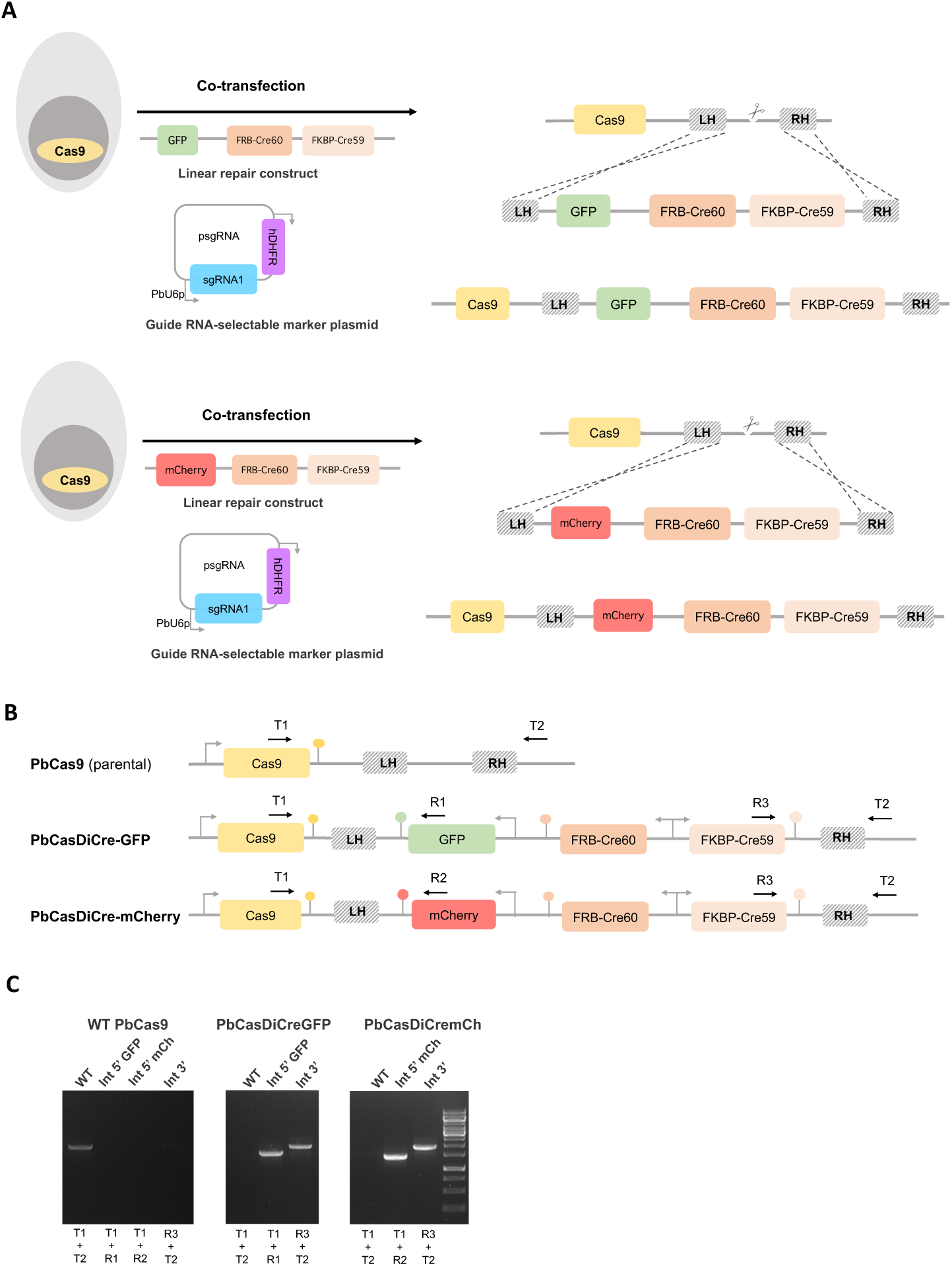
Generation of PbCasDiCre parasites. **A**. Strategy to integrate DiCre and GFP/mCherry cassettes into Cas9-expressing parasites using CRISPR. The PbCas9 parasites were co-transfected with a linearized DNA repair construct containing the two Cre components and a GFP or mCherry cassette, and with a plasmid encoding a sgRNA guide and a pyrimethamine-resistance cassette (hDHFR). Following Cas9-mediated DNA cleavage, the repair construct was integrated by double crossover recombination at the *p230p locus* downstream of the Cas9 cassette. LH, left homology arm; RH, right homology arm. **B.** Schematic of the *p230p* locus of parental PbCas9 and transgenic PbCasDiCre-GFP and PbCasDiCre-mCherry parasites. Genotyping primers are indicated by arrows (T, test; R, recombinant). **C.** PCR analysis of the genomic DNA obtained from the parental PbCas9 line and the two recombinant lines PbCasDiCre-GFP and PbCasDiCre-mCherry. Confirmation of the expected recombination events was achieved with primer combinations specific for 5’ or 3’ integration. A wild type-specific PCR reaction (WT) confirmed the absence of residual parental parasites in the PbCasDiCre-GFP and PbCasDiCre-mCherry transgenic lines.

Following the transfection of *PbCas9* parasites with the sgRNA plasmid and the linearized donor template DNA, integration of the repair construct by double homologous recombination results in the generation of parasites containing three cassettes integrated at the *p230p* locus: the Cas9, the DiCre and the GFP or mCherry cassettes (**Figure 1B**). Transfected parasites were exposed to pyrimethamine to select parasites containing the sgRNA plasmid. Following withdrawal of the selection drug, recombinant parasites were recovered and genotyped by PCR, using primer combinations specific for the parental or the recombined locus, respectively (**Figure 1C**). Genotyping PCR confirmed that the selected parasites had integrated the DiCre construct, with no remnant of the parental parasites (**Figure 1C**), consistent with 100% efficiency achieved with the integrated Cas9 approach [11]. PCR amplicons were verified by Sanger sequencing. We also conducted a genome wide sequence analysis using Oxford Nanopore Technology in order to verify the integrity of the DiCre and GFP/mCherry cassettes, and their correct integration at the expected locus. Alignment of the reads from PbCasDiCre-GFP and PbCasDiCre-mCherry parasite gDNA to the predicted recombined *p230p* locus showed an average coverage of approximately 60-70X encompassing the entire locus, confirming the correct integration of the fluorescent and DiCre cassettes downstream of Cas9, as expected (**Supplementary Figure S1**). In contrast, alignment of the reads from parental PbCas9 parasite gDNA to the predicted modified *p230p* locus of PbCasDiCre-GFP and PbCasDiCre-mCherry parasites showed 3 uncovered regions, corresponding to the GFP or mCherry cassette and the two DiCre components, respectively (**Supplementary Figure S1**). The recombinant parasite lines were exposed to 5-fluorocytosine (5-FC) to eliminate any residual parasite harboring the sgRNA plasmid, resulting in pure, drug-selectable marker free PbCasDiCre-GFP and PbCasDiCre-mCherry parasite populations.

### Transgenic PbCas-DiCre-GFP and PbCas-DiCre-mCherry parasites complete their life cycle normally

We examined the progression of the modified parasites through the life cycle, by comparing with lab-standardised control lines to exclude any possible effects of the introduction of the cassettes. Mice were infected with PbCasDiCre-GFP or PbCasDiCre-mCherry parasites and used to feed *A. stephensi* mosquitoes. We then monitored parasite development inside infected mosquitoes using fluorescence microscopy. Both transgenic lines colonized the mosquito midguts to form oocysts, and developed into sporozoites that invaded the mosquito salivary glands. Similar numbers of salivary gland sporozoites were obtained from mosquitoes infected with the modified parasite lines as compared to PbGFP or PbDiCre parasites used as controls (Figure 2A), showing that PbCasDiCre parasites develop normally inside the vector. To check the development of the mammalian liver stages, PbCasDiCre-GFP and PbCasDiCre-mCherry sporozoites were isolated from the salivary glands of infected mosquitoes and incubated with HepG2 cells. Transgenic sporozoites showed normal cell traversal activity, as determined using a dextran-based cell wound repair assay (Figure 2B), and formed similar numbers of exo-erythrocytic forms (EEFs) as the control PbGFP and PbDiCre parasites, as determined by flow cytometry 24 hours post-infection (Figure 2C) and by fluorescence microscopy 24 and 48 hours post-infection after labelling of the parasitophorous vacuole membrane marker UIS4 (Figure 2D**, 2E and 2F**). We also checked the infectivity of PbCasDiCre-GFP and PbCasDiCre-mCherry sporozoites *in vivo* in C57BL/6J mice. After intravenous inoculation of the parasites, mice injected with PbCasDiCre-GFP and PbCasDiCre-mCherry parasites all developed a patent parasitaemia, with similar pre-patency periods and blood-stage growth as control PbGFP parasites (Figure 2G). Altogether, these data confirm that *P. berghei* parasites constitutively expressing Cas9 and DiCre are capable of completing the parasite life cycle normally, and thus provide a suitable platform to study gene function across the life cycle.

**Figure 2.**
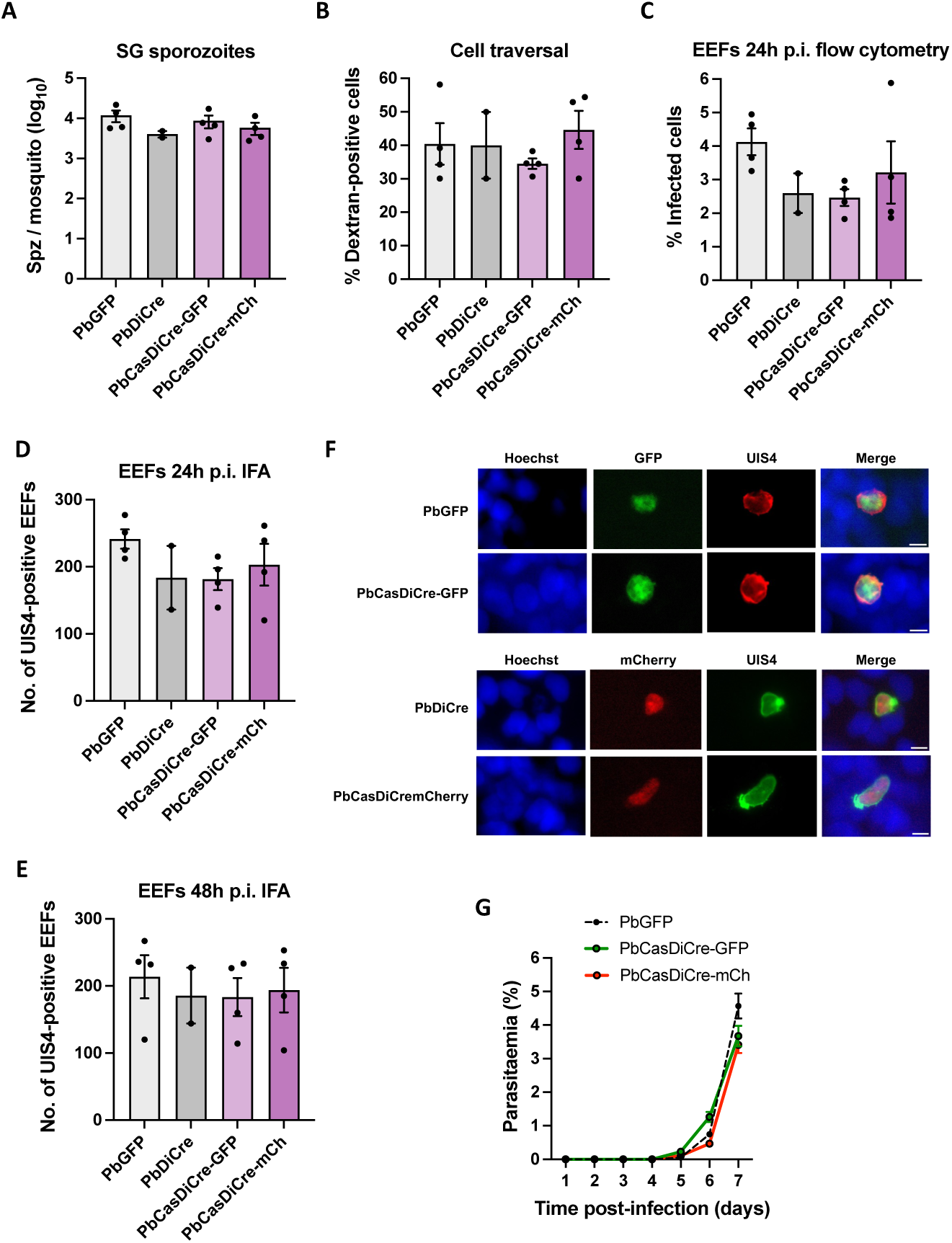
Phenotypic characterization of PbCasDiCre parasites. **A.** Comparison of sporozoite numbers isolated from salivary glands of female mosquitoes infected with the control lines, PbGFP and PbDiCre, or with the new PbCasDiCre-GFP and PbCasDiCre-mCherry transgenic lines. Results shown are mean +/- SEM of two to four independent experiments. **B.** Quantification of traversed (dextran-positive) HepG2 cells by FACS after incubation for 3h with PbGFP, PbDiCre, PbCasDiCre-GFP or PbCasDiCre-mCherry salivary gland sporozoites in the presence of fluorescent dextran. Results shown are mean +/- SEM of two to four independent experiments. **C**. Quantification of infected (GFP- or mCherry-positive) HepG2 cells by flow cytometry 24h post-infection with PbGFP, PbDiCre, PbCasDiCre-GFP or PbCasDiCre-mCherry salivary gland sporozoites. Results shown are mean +/- SEM of two to four independent experiments. **D-E**. Quantification of UIS4-labelled exo-erythrocytic forms (EEFs) in HepG2 cells as determined by fluorescence microscopy 24h (**D**) and 48h (**E**) post-infection. Results shown are mean +/- SEM of two to four independent experiments. **F.** Images of PbCasDiCre-GFP and PbCasDiCre-mCherry EEFs compared to the controls PbGFP and PbDiCre 48 hours p.i. in HepG2 cells. The parasitophorous vacuole membrane was labelled with anti-UIS4 antibodies and nuclei were stained with Hoechst 33342. Scale bar, 5 μm. **G.** Parasite development was compared *in vivo* post-sporozoite injection. C57BL6/J mice (n=3) were injected intravenously with 1 x 10^3^ PbGFP, PbCasDiCre-GFP or PbCasDiCre-mCherry sporozoites. Parasitaemia was then followed by flow cytometry. The data shown are mean +/- SEM of 3 mice per group.

### CRISPR-assisted conditional knockout of *clamp* in *P. berghei*

In order to validate the Cas9 and DiCre combination, we used the PbCasDiCre-GFP line to conditionally delete the gene encoding the Claudin-like apicomplexan microneme protein (CLAMP, PBANKA_0514200). CLAMP plays an essential role during invasion by *Toxoplasma gondii* tachyzoites and in *P. falciparum* asexual blood stages [13,14]. Being essential for blood stage growth, CLAMP is refractory to conventional gene deletion, thereby requiring a conditional genome editing strategy. In a previous study using PbDiCre parasites, we showed that CLAMP is involved during blood stage growth and in gliding motility and infectivity of sporozoites in *P. berghei* [8]. In that study, conditional knockout parasites were obtained after introducing LoxN sites upstream and downstream of *clamp* gene in a two-step strategy requiring two successive transfections [8]. The blood stage growth of CLAMP conditional knockout parasites exposed to rapamycin was reduced as compared to control parasites. Here, in order to allow functional studies in merozoites, we used a different one-step strategy to introduce a first LoxP site inside an artificial intron, as described in *P. falciparum* [5], and a second LoxP site immediately downstream of the STOP codon (Figure 3A). A triple HA tag was introduced before the STOP codon for C-terminal tagging of CLAMP. Using this strategy, CRE-mediated recombination should result in immediate and complete suppression of CLAMP expression. Two guide RNAs were selected in the *clamp* locus, in the 5’ and 3’ regions of the coding sequence, respectively, and cloned into a single plasmid containing PbU6 and PfU6 promoters (Figure 3A) [11]. The donor DNA template carrying a modified *clamp* gene with an artificial intron taken from the *P. berghei slarp* gene (PBANKA_0902100) [15] and containing the first LoxP site, a 3xHA tag and a second LoxP site, was provided as a synthetic gene and linearized before transfection.

**Figure 3.**
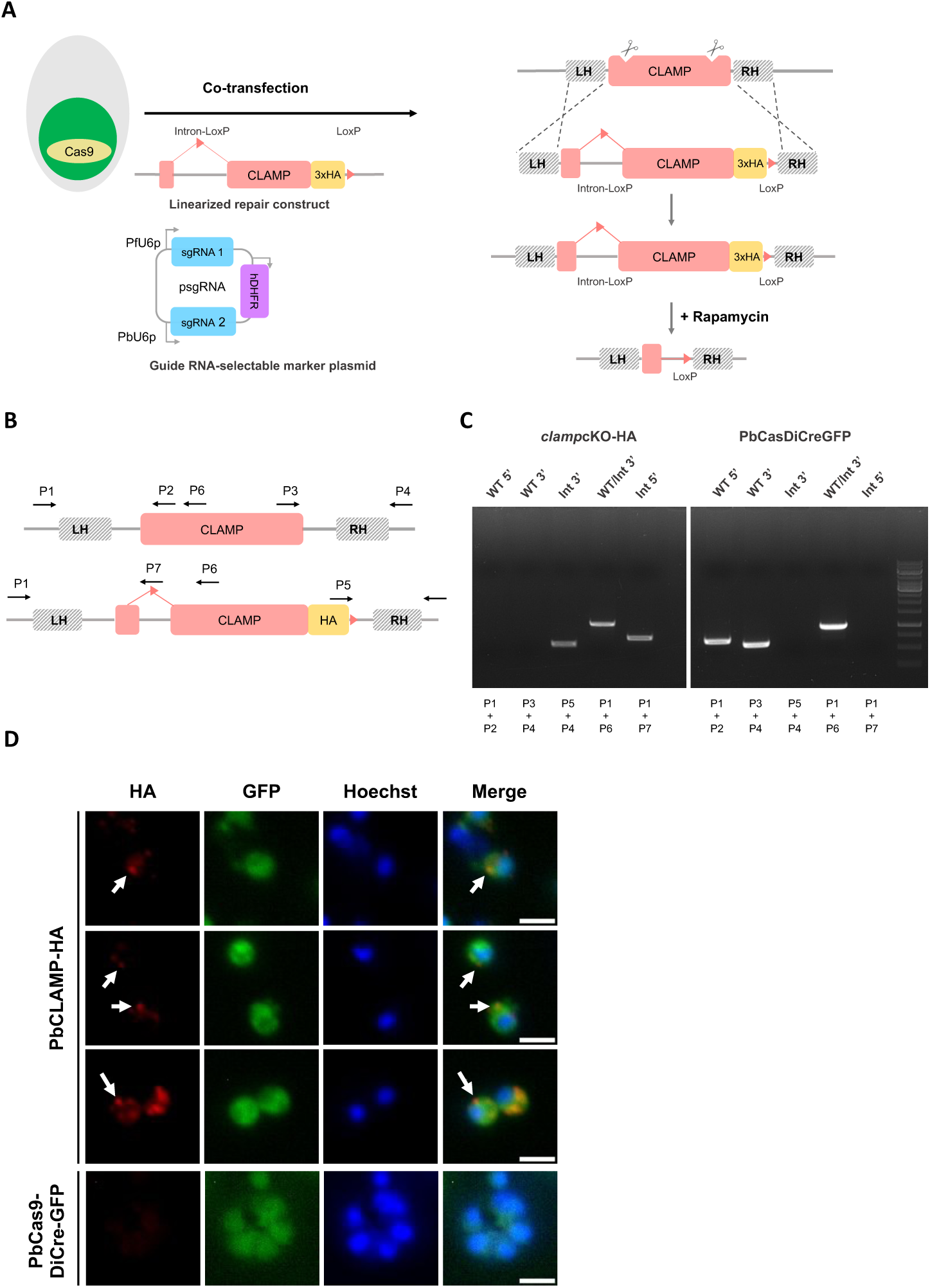
Generation of *clamp*cKO-HA parasites. **A.** Strategy to flox the *clamp* gene in PbCasDiCre-GFP using CRISPR. PbCasDiCre-GFP parasites were co-transfected with a linearized DNA repair construct containing a floxed *clamp* gene with a C-terminal 3xHA tag, and with a plasmid encoding two sgRNA guides and a pyrimethamine-resistance cassette (hDHFR). Following Cas9-mediated DNA cleavage, the repair construct was integrated by double crossover recombination at the *clamp* locus, resulting in the generation of the *clamp*cKO-HA parasite line. **B.** Schematic of the *clamp* locus of parental and transgenic *clamp*cKO-HA parasites. Genotyping primers are indicated by arrows. **C.** PCR analysis of the genomic DNA obtained from the parental PbCasDiCre-GFP and the recombinant *clamp*cKO-HA lines. Confirmation of the expected recombination events was achieved with primer combinations specific for 5’ or 3’ integration. Wild type-specific PCR reactions (WT 5’ and 3’) confirmed the absence of residual parental parasites in the *clamp*cKO-HA line. **D.** Immunofluorescence analysis of *clamp*cKO-HA merozoites using anti-HA antibodies revealed a polar and punctate distribution of the protein, with no labelling observed in parental untagged parasites. Scale bar, 2 μm.

Following the transfection of PbCasDiCre-GFP parasites with the double sgRNA plasmid targeting *clamp* and the linearized donor template DNA, integration of the repair construct by double homologous recombination results in the generation of parasites containing a floxed *clamp* gene with a C-terminal 3xHA tag (Figure 3A). Recombinant parasites were selected with pyrimethamine and genotyped by PCR, using primer combinations specific for the parental or the recombined locus, respectively (Figure 3B-C). Genotyping PCR confirmed that the selected parasites had integrated the repair construct, with no remnant of the parental parasites (Figure 3C). PCR amplicons were verified by Sanger sequencing. We also conducted a genome wide sequence analysis using Oxford Nanopore Technology in order to verify the integrity of the modified *clamp* locus in the *clamp*cKO-HA line. Reads from the *clamp*cKO parasite DNA aligned entirely to the expected modified *clamp*cKO locus on chromosome 5, while alignment of the reads from parental PbCasDiCre-GFP showed 2 uncovered regions around the *clamp* gene locus, matching the LoxP and 3xHA/LoxP sequences, respectively (**Supplementary Figure S2**).

Analysis of *clamp*cKO-HA merozoites by immunofluorescence using anti-HA antibodies revealed a punctate distribution of the protein which was often predominant at one pole of the parasite (Figure 3D), reminiscent of the apical accumulation of the protein previously seen in sporozoites [8]. As a control, no labelling was observed in untagged parasites (Figure 3D).

We then assessed the effects of rapamycin on *clamp*cKO parasites during blood-stage growth. A first group of mice with patent parasitaemia was treated with a single oral dose of rapamycin, while mice in a control group were left untreated. We then monitored the parasitaemia over time by flow cytometry. Remarkably, rapamycin-induced excision of *clamp* reduced parasitaemia to nearly zero in a single cycle (<24 hours), as compared to untreated parasites (Figure 4A), in agreement with an essential role for CLAMP in asexual blood stages. In order to verify that *clamp* gene excision induced by rapamycin exposure was efficient in depleting CLAMP protein, we exposed *clamp*cKO parasites to rapamycin *in vitro* and analyzed the abundance of CLAMP by Western blot using anti-HA antibodies. While in untreated parasites the protein was detected as a single ∼40 kDa band, consistent with data obtained previously with sporozoites expressing Flag-tagged CLAMP [8], CLAMP was no longer detected in merozoites after rapamycin treatment (Figure 4B). The rapid and dramatic decrease of parasitaemia during the first cycle after exposure to rapamycin suggests that CLAMP-deficient parasites are not able to invade erythrocytes. To directly assess the invasive capacity of CLAMP-deficient merozoites, *clamp*cKO merozoites were produced in culture in the presence or absence of rapamycin, and injected intravenously into mice. Following inoculation, untreated merozoites efficiently invaded mouse erythrocytes and established a blood stage infection, as evidenced by flow cytometry (Figure 4C). In sharp contrast, rapamycin-exposed merozoites failed to invade erythrocytes, as evidenced by the absence of detectable parasitaemia after inoculation into mice (Figures 4C). These data thus demonstrate that CLAMP is required in *P. berghei* merozoites for invasion of erythrocytes, and illustrate the efficacy of the combined Cas9-DiCre approach to investigate parasite gene function.

**Figure 4.**
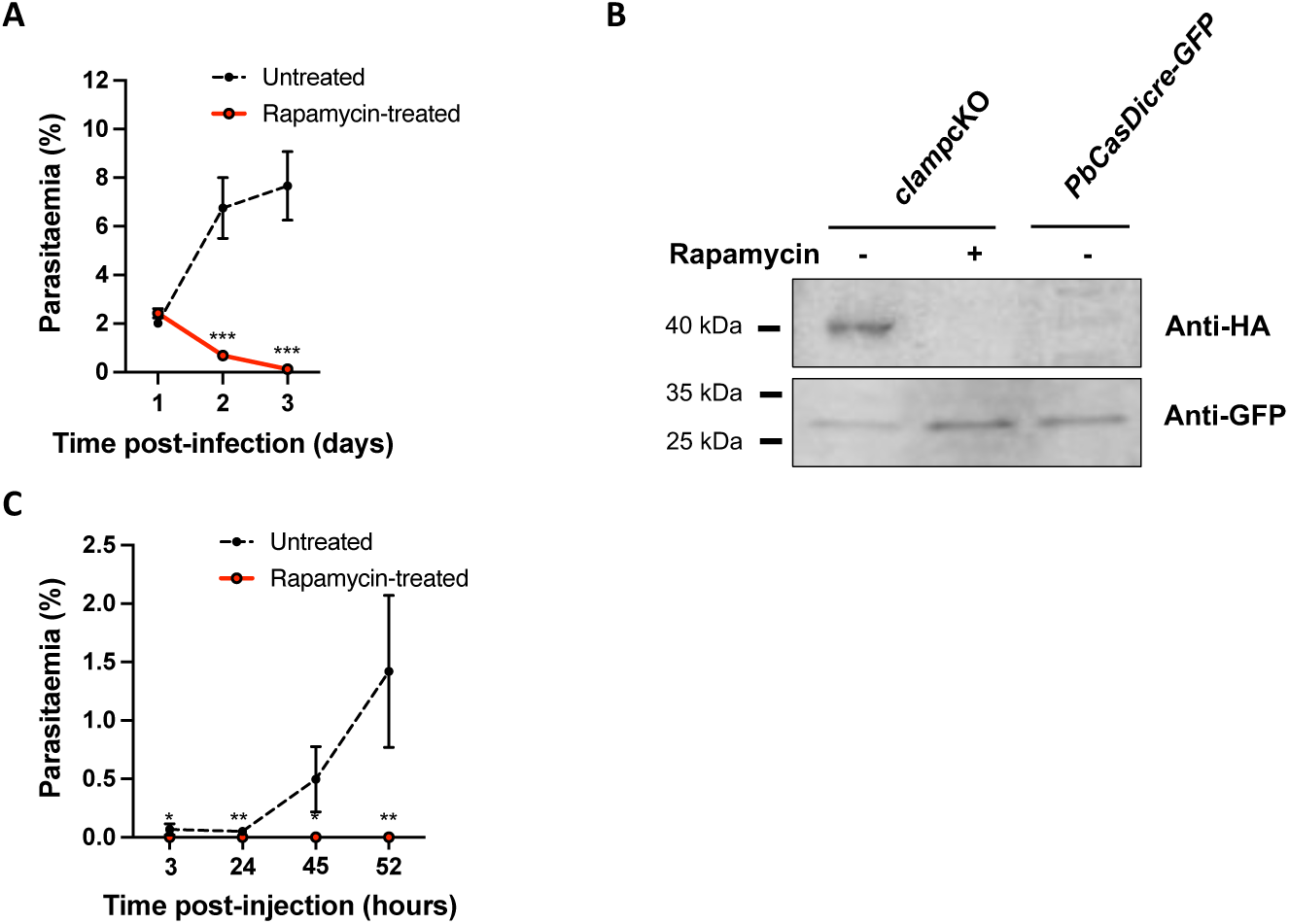
Conditional knockout of *clamp* abrogates merozoite infectivity. **A.** Blood stage growth of rapamycin-treated and untreated *clamp*cKO-HA parasites. Rapamycin was administered at day 1. The graph shows the parasitaemia (mean +/- SEM) in groups of 5 mice, as quantified by flow cytometry based on GFP detection. ***, p<0.001 (unpaired t test). **B.** Western blot analysis of blood stage schizont lysates from untreated or rapamycin-treated *clamp*cKO-HA parasites, and control parental PbCasDiCre-GFP parasites, using anti-HA antibodies to detect CLAMP. GFP was used as a loading control. The uncropped images of the blots are shown in Supplementary Figure S3. **C.** *clamp*cKO-HA parasites were cultured for 18h in the presence or absence of rapamycin. Merozoites were then purified and injected intravenously into mice. The graph shows the parasitaemia (mean +/- SEM) in groups of 3 mice, as quantified by flow cytometry based on GFP detection. *, p<0.05; **, p<0.01 (unpaired t test).

## Discussion

CRISPR has become a standard method for genetic modification in many organisms, including *Plasmodium* spp and other apicomplexan parasites [16]. In the rodent malaria model parasite *P. berghei*, improvement of the CRISPR/Cas9 system has been recently achieved through the constitutive expression of Cas9 and the usage of a linear donor template, allowing robust and highly accurate genome editing [11]. Following transfection of Cas9-expressing parasites with a sgRNA-encoding plasmid, site specific DSB introduced in the genome by Cas9 result in parasite death, unless DNA is repaired by homologous recombination with the linear donor DNA template provided with the sgRNA plasmid during transfection. A short drug selection step eliminates parasites that have not taken up the sgRNA plasmid. This strategy allows recovering recombinant parasites with very high efficiency, close to 100%. Here we used this strategy to modify by CRISPR the dispensable *p230p* locus in PbCas parasites to introduce an additional DiCre cassette in the genome of PbCas9 parasites, together with a fluorescence GFP or mCherry cassette to facilitate parasite monitoring. Following transfection, we recovered pure populations of recombinant parasites expressing the combined cassettes, with no detectable persistence of the parental parasites, illustrating the very high efficiency of the CRISPR approach. We show here that Cas-DiCre parasites progress normally through the life cycle, in both the mosquito and mammalian hosts, confirming that expression of Cas9 and DiCre components has no deleterious effect on the parasite viability and infectivity, as previously observed when each component was integrated independently in the parasite genome [7,11].

We have previously implemented the DiCre system in *P. berghei* parasites and illustrated how this approach can be used to conditionally delete essential genes before transmission to mosquitoes to investigate gene function in sporozoites [7–9,17]. This strategy allowed us to demonstrate the role of AMA1 and RONs during invasion of mosquito salivary glands and mammalian hepatocytes [9], as well as the role of CLAMP in sporozoite motility and infectivity [8]. In these studies, Lox sites were inserted upstream and downstream of the target genes in two steps, requiring two successive transfections. Another limitation of the standalone DiCre approach was that a drug resistance cassette had to be integrated for the selection of modified parasites, thus limiting iterative genetic modifications. Here, we used CRISPR to insert two Lox sites in *clamp* gene in one single transfection. A short selection step with pyrimethamine was only used to select for parasites harboring the sgRNA plasmid, and following drug withdrawal we obtained pure populations of drug selectable marker-free *clamp*cKO parasites. In addition, one of the Lox sites was inserted inside a synthetic intron, as previously described in *P. falciparum* [5]. The advantage of such a strategy is that rapamycin-induced recombination results in immediate disruption of the expression unit, allowing phenotypic analysis in merozoites.

We have previously shown that CLAMP is required for *P. berghei* blood stage growth [8]. Here, rapamycin-induced disruption of *clamp* did not impair maturation of schizonts and production of merozoites *in vitro*. However, CLAMP-deficient merozoites were not capable of reinvading erythrocytes *in vivo* after inoculation into mice. This result demonstrates that CLAMP is required for host cell invasion, as described in *Toxoplasma gondii* [13,14]. Immunofluorescence assays revealed a punctate distribution of HA-tagged CLAMP in merozoites. Whether CLAMP localizes to specific subsets of micronemes in merozoites and contributes to rhoptry secretion during host cell invasion, as in *T. gondii* tachyzoites [13,14], deserves further investigation.

This study illustrates that the combination of Cas9 and DiCre systems provides a robust and versatile platform allowing diverse genome editing strategies in *P. berghei*, including conditional gene disruption to decipher the function of essential genes across the parasite life cycle. New CRISPR-based methods are emerging in *Plasmodium*, including scalable genetic screens based on Cas9, as recently implemented in *P. falciparum* [18] and *P. berghei* [19]. The Cas-DiCre *P. berghei* lines may represent valuable tools for such CRISPR-based genetic screens, enriching our toolbox for functional studies in the malaria parasite.

## Materials and methods

### Ethics Statement

All animal work was conducted in strict accordance with the Directive 2010/63/EU of the European Parliament and Council on the protection of animals used for scientific purposes. Protocols were approved by the Ethics Committee Charles Darwin N°005 (approval #24845-2020032816248463).

### Generation of plasmids for transfection

*sgRNA guide plasmids*.

Using the Chop-Chop (https://chopchop.cbu.uib.no) and Benchling (https://www.benchling.com) programs, one or two 19-20 bp guide RNA sequences were selected upstream of PAM motifs in each of the target genes (*p230p* or *clamp*). Their complementary oligonucleotides were subsequently designed and optimized in the Takara Primer design tool (https://www.takarabio.com/learning-centers/cloning/primer-design-and-other-tools). A guanosine nucleotide was added at the 5’ end of the forward oligonucleotide for enhancing transcriptional initiation [11]. Paired oligonucleotides were annealed and cloned into a *Bsm*BI- or *Bsa*I-digested psgRNA_Pf-Pb U6_2targets plasmid using the In-Fusion HD Cloning Kit (Takara), resulting in the insertion of the guide RNA immediately downstream of PfU6 or PbU6 promoter, respectively [11]. This plasmid contains a *hDHFR*-*yfcu* cassette, for positive selection by pyrimethamine and negative selection by 5-fluorocytosine (5-FC) [20,21]. The resulting sgRNA guide plasmids were checked by Sanger sequencing prior to transfection.

*Donor DNA templates for DNA repair by double homologous recombination*.

Donor DNA templates for integrating the DiCre and GFP/mCherry cassettes were assembled as plasmids in four steps. First, the entire DiCre cassette, containing the Cre59 and Cre60 fragments under control of the bidirectional eEF1α promoter and followed by 3’ UTR sequences of PfCAM and PbHSP70, respectively, was amplified by PCR using the previously described DiCre construct [7] as a template, and cloned into a pBluescript backbone between *Sal*I and *Eco*RI sites. Then, fragments from *p230p* corresponding to 5’ and 3’ homology regions were cloned on each side of the DiCre cassette, between *Kpn*I and *Xho*I sites or *Not*I and *Sac*II sites, respectively. Finally, a fluorescence expression cassette, consisting of either GFP or mCherry under control of *pbhsp70* promoter and the 3’ UTR of *pbdhfr*, was amplified by PCR using genomic DNA from either PbGFP or PbDiCre parasites, respectively, and inserted in the construct at the *Xho*I site. All cloning steps were performed using the CloneAmp HiFi PCR premix and the In-Fusion HD Cloning Kit (Takara). The resulting plasmids were verified by Sanger sequencing (Eurofins Genomics) and linearized with *Nhe*I and *Sac*II before transfection. The donor template to modify *clamp* locus was provided as a synthetic gene, containing a 529-bp 5’ upstream fragment immediately upstream of the ATG of *clamp*, serving for 5’ homologous recombination, a 31-bp sequence of the ORF, a 170-bp intron from *slarp* gene containing a LoxP site, a 2643-bp fragment corresponding to the rest of the ORF, a triple HA epitope tag, a STOP codon, a second LoxP site, and a 528-bp 3’ downstream fragment serving for 3’ homologous recombination. The entire synthetic gene was flanked by two *Xho*I sites allowing linearization prior to transfection. All the primer sequences are listed in **Table S1**.

### Experimental animals, parasites and cell lines

Female SWISS mice (6–8 weeks old, from Janvier Labs) were used for all blood stage parasite infections. Parasite lines were maintained in mice through intraperitoneal injections of parasite cryostocks and transmitted to *Anopheles stephensi* mosquitoes for experimental purposes. *P*. *berghei* sporozoites were isolated from infected female *Anopheles* mosquitoes and intravenously injected into C57BL/6J mice (female, 4–6 weeks old, Janvier Labs). A drop of blood from the tail was collected in 1ml PBS daily and used to monitor the parasitaemia by flow cytometry. For parasite transfection, schizonts were purified from an overnight culture of the parent parasite line PbCas9 [11] and transfected with a mix of 10 μg of sgRNA plasmid and 10 μg of linearized donor template by electroporation using the AMAXA Nucleofector device (program U033), as previously described [22], and immediately injected intravenously into the tail vein of SWISS mice. To permit the selection of resistant transgenic parasites, pyrimethamine (35 mg/L) was added to the drinking water and administered to mice, starting one day after transfection and for a total of 4-5 days. Following withdrawal of pyrimethamine, the mice were monitored daily by flow cytometry to detect the reappearance of parasites. When parasitaemia reached at least 1%, mouse blood was collected for preparation of cryostocks and isolation of parasites for genomic DNA extraction and genotyping. To eliminate parasites carrying the sgRNA plasmid, an additional step of negative selection was performed, by exposing infected mice to 5-FC (Meda Pharma) at 1 mg/mL in the drinking water.

*Anopheles stephensi* mosquitoes were reared at 24–26°C with 80% humidity and permitted to feed on infected mice that were anaesthetized, using standard methods of mosquito infection as previously described [23]. Post-feeding, *P*. *berghei*-infected mosquitoes were kept at 21°C and fed on a 10% sucrose solution. Salivary gland sporozoites were collected from infected mosquitoes between 21 and 28 days post-feeding, by hand dissection and homogenisation of isolated salivary glands in complete DMEM (DMEM supplemented with 10% FCS, 1% Penicillin-Streptomycin and 1% L-Glutamine). Mosquitoes infected with GFP- or mCherry-expressing parasites were sorted under a fluorescence microscope prior to dissection. HepG2 cells (ATCC HB-8065) were cultured in complete DMEM, as previously described [24].

### Rapamycin treatment

DiCre recombinase-mediated excision of targeted DNA sequences was achieved by administration of a single dose of 200 μg Rapamycin (1mg/ml stock, Rapamune, Pfizer) to mice by oral gavage, or by culturing infected erythrocytes for 18 hours in the presence of 100 nM rapamycin (from 1 mM stock solution in DMSO, Sigma-Aldrich). Control cultures were exposed to DMSO alone (0.01% vol/vol).

### Genotyping PCR

Blood collected from infected mice was passed through a CF11 column (Whatman) to deplete leucocytes. The collected RBCs were then centrifuged and lysed with 0.2% saponin (Sigma) to recover parasite material for genomic DNA isolation using a kit (Qiagen DNA Easy Blood and Tissue Kit), according to the manufacturer’s instructions. Genomic DNA served as template for PCR, using specific primer combinations designed to detect the wild-type or recombined loci. All PCR reactions were carried out using Recombinant Taq DNA Polymerase (5U/μL from Thermo Scientific) and standard PCR cycling conditions. All the primer sequences are listed in **Table S1**.

### Oxford Nanopore Technology sequencing

#### DNA sample preparation

Whole blood was collected from two infected mice at 5-10% parasitaemia and depleted from white blood cells by successive passage on a CF11 column (Whatman) and a Plasmodipur filter (R-Biopharm). After centrifugation of the filtered blood at 1500 rpm for 8 min at room temperature, the supernatant was discarded and the RBC pellet was resuspended in 0.2% saponin (Sigma) in PBS. Parasites were then pelleted by centrifugation at 2800 rpm for 8 min at room temperature and lysed for DNA extraction using the genomic DNA Easy Blood and Tissue kit (Qiagen, cat. #69504), according to the manufacturer’s instructions. DNA was kept at 4°C and never frozen. Following DNA purification, the sample volume was brought to 200 μL with the elution buffer and gDNA was extracted twice with 200 μL Phenol/Chloroform/Isoamyl alcohol to eliminate hemozoin contamination. The total aqueous fraction (360 μL) was treated with the same volume of Chloroform and the top aqueous phase was collected after centrifugation at maximal speed for 5 min at 4°C. gDNA was supplemented with 1 μL glycogen (10 mg/mL) and was precipitated with 1/10 volume of Sodium Acetate 3M pH 5.5 and 2 volumes of 100% ethanol. After overnight incubation at −20°C, gDNA was centrifuged at maximal speed for 20 min at 4°C, washed twice with 75% ethanol and air-dried for 5 min at room temperature.

#### Nanopore library preparation and sequencing

The quality, quantity and size of gDNAs from each strain were verified by spectrophotometry (Nanodrop), fluorimetry (Qubit) and microelectrophoresis (TapeStation 4151), respectively, and correspond to the recommendations of Oxford Nanopore Technologies. Two pools of 6 libraries were prepared from at least 400 ng of gDNA from each parasite strain using the SQK-NBD114-24 ligation kit following the supplier’s protocol. The gDNAs from one PbCas9, two PbCasDiCre-GFP, two PbCasDiCre-mCherry and one *clamp*cKO sample were multiplexed in each pool. The first pool was sequenced on a MinION R10.4.1 flow cell (Cat#FLO-MIN114) for 30h, then the flow cell was washed with the kit (WSH004) following the manufacturer’s protocol and the second pool was sequenced for 66h on the same flow cell. Fast5 raw data was generated using ONT’s MinKnow v 23.04.5 software and demultiplexing was performed in real time using guppy software (version 6.5.7).

#### Preparation of reference genomes

The fasta files used as reference genome for read alignment for the PbCas, PbCasDiCre-GFP, PbCasDiCre-mCherry strains were generated by inserting the sequence of the Cas9, DiCre-GFP and DiCre-mCherry cassettes, respectively, in place of the sequence between bases 196,621 to 213,484 of chromosome 3 of the genome of *P. berghei* strain ANKA ‘’PlasmoDB-64_PbergheiANKA_Genome’’ from the PlasmoDB database. The reference genome of the *clamp*cKO strain was generated from the fasta file of the PbCasDiCre-GFP strain by inserting the floxed *clamp* cassette into chromosome 5 in place of the sequence between bases 534,064 and 544,607.

#### Fastq analysis

Fastq files were generated in Super Accuracy (SupAC) mode from Fast5 files using the dorado basecaller (v7.3.11) of the Minknow software (v24.02.19). Fastq files were analyzed on the Galaxy Nanopore Europe server with the tools Porechop (v.0.2.4) for read trimming and map with minimap2 (v2.28) for alignments. Reads from PbCas (parental), PbCasDiCre-GFP and PbCasDiCre-mCherry DNA were aligned to the PbCasDiCre-GFP or PbCasDiCre-mCherry reference genomes, respectively. Reads from the PbCasDiCre-GFP (parental) and *clamp*cKO DNA were aligned on the reference *clamp*cKO genome. Alignment files were merged with the Merge BAM Files tool (v1.2.0) for each strain. Coverage was visualized in igv (v2.16.1).

### *In vitro* infections

HepG2 cells were seeded in collagen-coated culture plates, at a density of 30,000 cells/well in a 96-well plate for flow cytometry analysis and immunofluorescence assays, 24 hours prior to infection with sporozoites. On the day of infection, the culture medium in the wells was refreshed with complete DMEM, followed by the addition of 10,000 sporozoites/well for flow cytometry analysis or 1,000 sporozoites/well for immunofluorescence assays. Infected cultures were incubated for 3 hours at 37°C. The wells were then washed twice with complete DMEM and then incubated for another 24-48 hours at 37°C before analysis by flow cytometry or fixation for immunofluorescence. For quantification of infected cells by flow cytometry, the cultures were trypsinized after two washes with PBS, followed by addition of complete DMEM and one round of centrifugation. After discarding the supernatant, the cells were directly resuspended in FACS buffer (PBS + 1% FCS) and analyzed on a Guava EasyCyte 6/2L bench cytometer equipped with 488 nm and 532 nm lasers (Millipore). Cell traversal activity was monitored by flow cytometry using a membrane wound-repair assay based on uptake of fluorescent dextran [25]. For this purpose, HepG2 cells were incubated for 3 hours at 37°C with PbCasDiCre-GFP or PbCasDiCre-mCherry sporozoites in the presence of 1mg/mL rhodamine- or FITC-conjugated dextran, respectively, before analysis by flow cytometry.

### Immunofluorescence assays

For immunofluorescence assays on HepG2 infected cultures, the cells were washed twice with PBS, then fixed with 4% PFA for 10 minutes followed by two washes with PBS, quenching with 0.1 M glycine for 5 minutes, permeabilization with 1% Triton X-100 for 5 minutes before washes with PBS and blocking in PBS with 3% bovine serum albumin (BSA). Cells were then incubated for 1h with goat anti-UIS4 primary antibody (1:500, Sicgen), followed by Alexa Fluor 594- or Alexa Fluor 488-conjugated donkey anti-goat secondary antibodies (1:1000, Life Technologies). Nuclei were stained with Hoechst 33342 (Life Technologies). Samples were then imaged on a Zeiss Axio Observer.Z1 fluorescence microscope equipped with a LD Plan-Neofluar 40x/0.6 Corr Ph2 M27 objective. The same exposure conditions were maintained for all the conditions in order to allow comparisons. Images were processed with ImageJ for adjustment of contrast. For immunofluorescence on infected erythrocytes, cultures of *clamp*cKO-HA and parental (PbCasDiCre-GFP) schizonts were fixed in 4% PFA as described above and permeabilized with Triton X-100. The HA tag was revealed using a rat monoclonal antibody (3F10, Roche) and Alexa Fluor 594-conjugated goat anti-rat secondary antibodies (1:1000, Life Technologies).

### Western blot

Mouse blood infected with *clamp*cKO-HA parasites (or PbCasDiCre-GFP as a negative control) was cultured overnight in the presence or absence of rapamycin, collected and resuspended in 1X PBS. Parasite pellets were then isolated by centrifugation, resuspended in Laemmli buffer and analyzed by SDS-PAGE under non-reducing conditions. Western blotting was performed using a rat monoclonal antibody against HA (Roche, 3F10) or a control goat polyclonal antibody against GFP (Sicgen, AB0066-20), followed by secondary antibodies coupled with Alexa Fluor 680 or 800 against rat or goat, respectively. Membranes were then analyzed using the InfraRed Odyssey system (Licor).

### Statistical analysis

Statistical significance was assessed by one-way or two-way ANOVA or unpaired t tests, as indicated in the figure legends and **Table S2**. All statistical tests were computed with GraphPad Prism 10 (GraphPad Software). *In vitro* experiments were performed with a minimum of three technical replicates per experiment. Quantitative source data are provided in **Table S2**.

## Acknowledgements

We thank Jean-François Franetich and Thierry Houpert for rearing of mosquitoes. This work was funded by grants from the Laboratoire d’Excellence ParaFrap (ANR-11-LABX-0024), the Agence Nationale de la Recherche (ANR-20-CE18-0013) and the Fondation pour la Recherche Médicale (EQU201903007823). This work benefited from equipment and services from the P3S core facility, a platform supported by the Conseil Régional d’Ile-de-France, Sorbonne Université, the National Institute for Health and Medical Research (Inserm) and the Biology, Health and Agronomy Infrastructure (IBiSA).

---

**Supplementary Figure S1.**
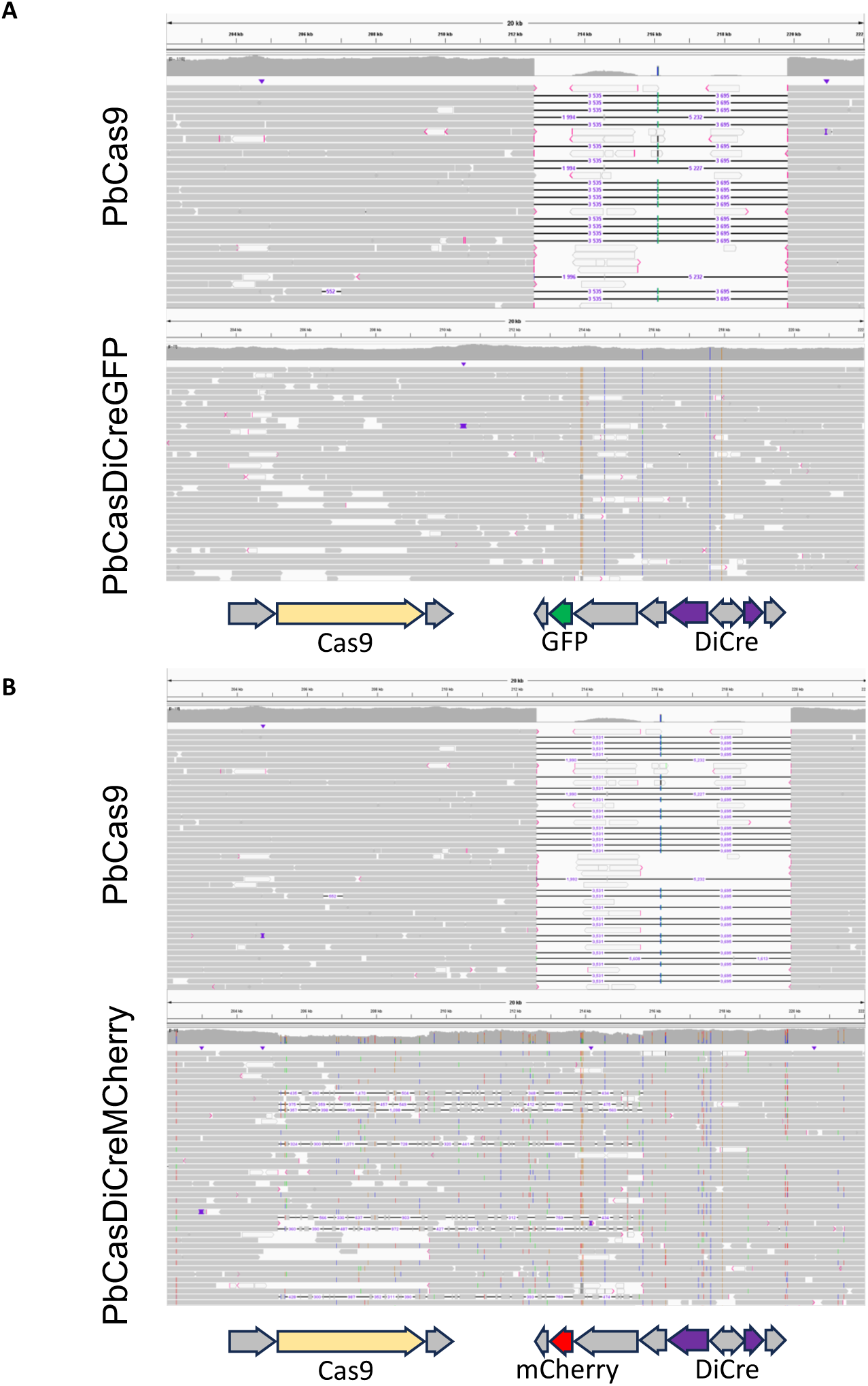
Nanopore sequencing of PbCas9, PbCas-DiCre-GFP and PbCas-DiCre-mCherry parasite lines Genomic DNA from PbCas9, PbCasDiCre-GFP and PbCasDiCre-mCherry parasites was sequenced by Oxford Nanopore Technology. Reads were aligned to the expected recombined genome sequences of PbCasDiCre-GFP (**A**) or PbCasDiCre-mCherry (**B**) parasites. Only the *p230p* locus on chromosome 3 is shown.

**Supplementary Figure S2.**
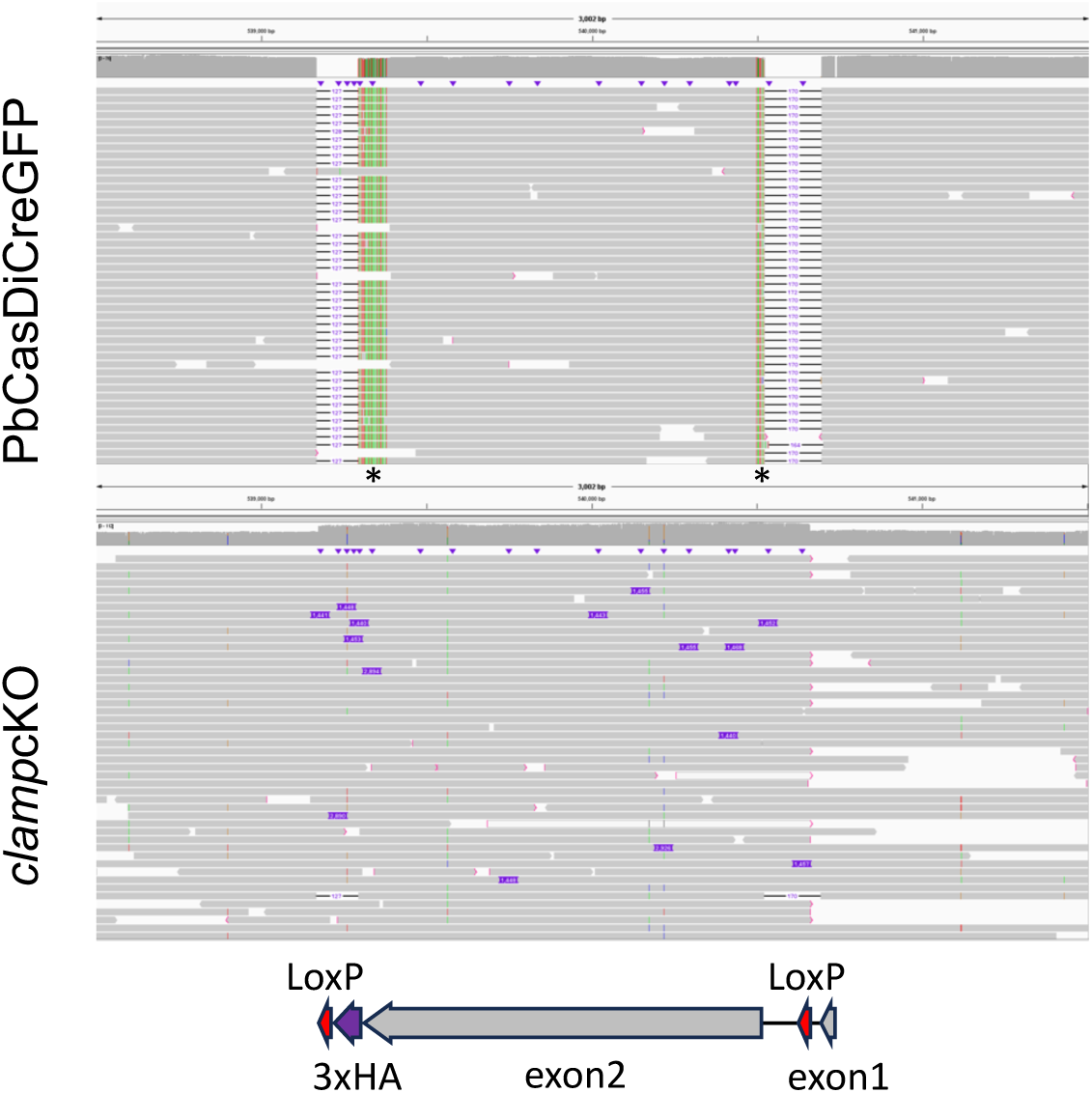
Nanopore sequencing of PbCas-DiCre-GFP and *clamp*cKO-HA parasite lines Genomic DNA from PbCasDiCre-GFP and *clamp*cKO-HA parasites was sequenced by Oxford Nanopore Technology. Reads were aligned to the expected recombined genome sequence. Only the *clamp* locus on chromosome 5 is shown.

**Supplementary Figure S3.**
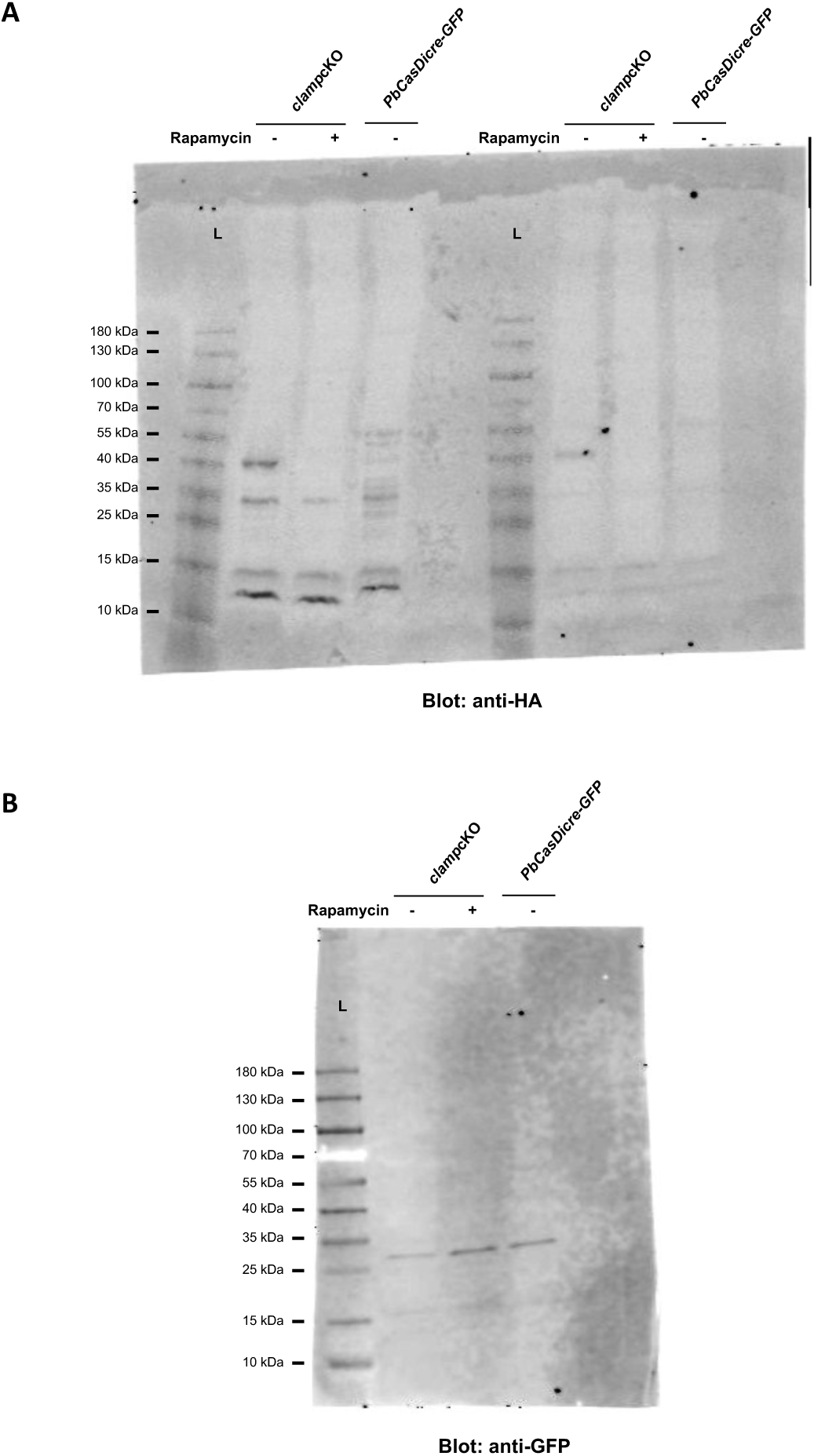
Uncropped blot images The uncropped blot images correspond to Figure 4C.

